# A novel glycosyltransferase organizes fungal glycogen and cell wall glucans

**DOI:** 10.1101/2023.10.24.563850

**Authors:** Liza Loza, Tamara L. Doering

## Abstract

Glycogen is a glucose storage molecule comprised of branched α-1,4-glucan chains, best known as an energy reserve that can be broken down to fuel central metabolism. Because fungal cells have a specialized need for glucose in building cell wall glucans, we investigated whether glycogen is used for this process. For these studies we focused on the pathogenic yeast *Cryptococcus neoformans*, which causes over 112,000 deaths per year worldwide. We identified two enzymes that influence formation of both glycogen and the cell wall: Gly-cogenin (Glg1), which initiates glycogen synthesis, and a novel protein that we call **<u>G</u>**lucan **<u>o</u>**rganizing **<u>e</u>**nzyme 1 (Goe1). We discovered that cells missing Glg1 lack α-1,4-glucan in their walls, indicating that this material is derived from glycogen. Without Goe1, glycogen rosettes are mislocalized and β-1,3-glucan in the cell wall is reduced. Altogether, our results provide mechanisms for a close association between glycogen and cell wall.

## INTRODUCTION

Storing carbohydrates as polymers is an ancient metabolic strategy conserved across all domains of life. One such polymer, glycogen, can be broken down in times of carbon scarcity. Glycogen features a protein core, called glycogenin, which is surrounded by chains of α-1,4-linked glucose (Glc) with α-1,6-linked branchpoints every 8-13 Glc units (depending on the organism)^1^. Approximately seven tiers of these branches radiate compactly from glycogenin, forming a dense 20-25 nm granule with a molecular weight of 10^4^-10^5^ kD^2,3^.

Glycogen synthesis and degradation were first glimpsed as biochemical activities present in muscle extracts and later mapped out in the model yeast *Saccharomyces cerevisiae* (*Sc*). In *Sc*, glycogenin proteins (Glg1 and Glg2) first catalyze the transfer of Glc from uridine diphosphate glucose (UDP-Glc) to internal tyro-sine (Tyr) residues, in an unusual self-glycosylation reaction^4,5^ (Figure 1A, green boxes). In the next phase of synthesis, glycogenin primes the chain by adding up to 10 α-1,4-linked Glc units to the Glc 1-*O*-tyrosyl group. Next, glycogen synthases (Gsy1 and Gsy2) extend this primer by processively adding α-1,4-linked Glc units, again using UDP-Glc as a reaction donor^6,7^. Finally, branchpoints are generated every 8-10 units when a branching enzyme (Glc3) transfers a distal block of α-1,4-linked Glc units to an interior Glc, forming an α-1,6 linkage^8,9^.

**Figure 1.**
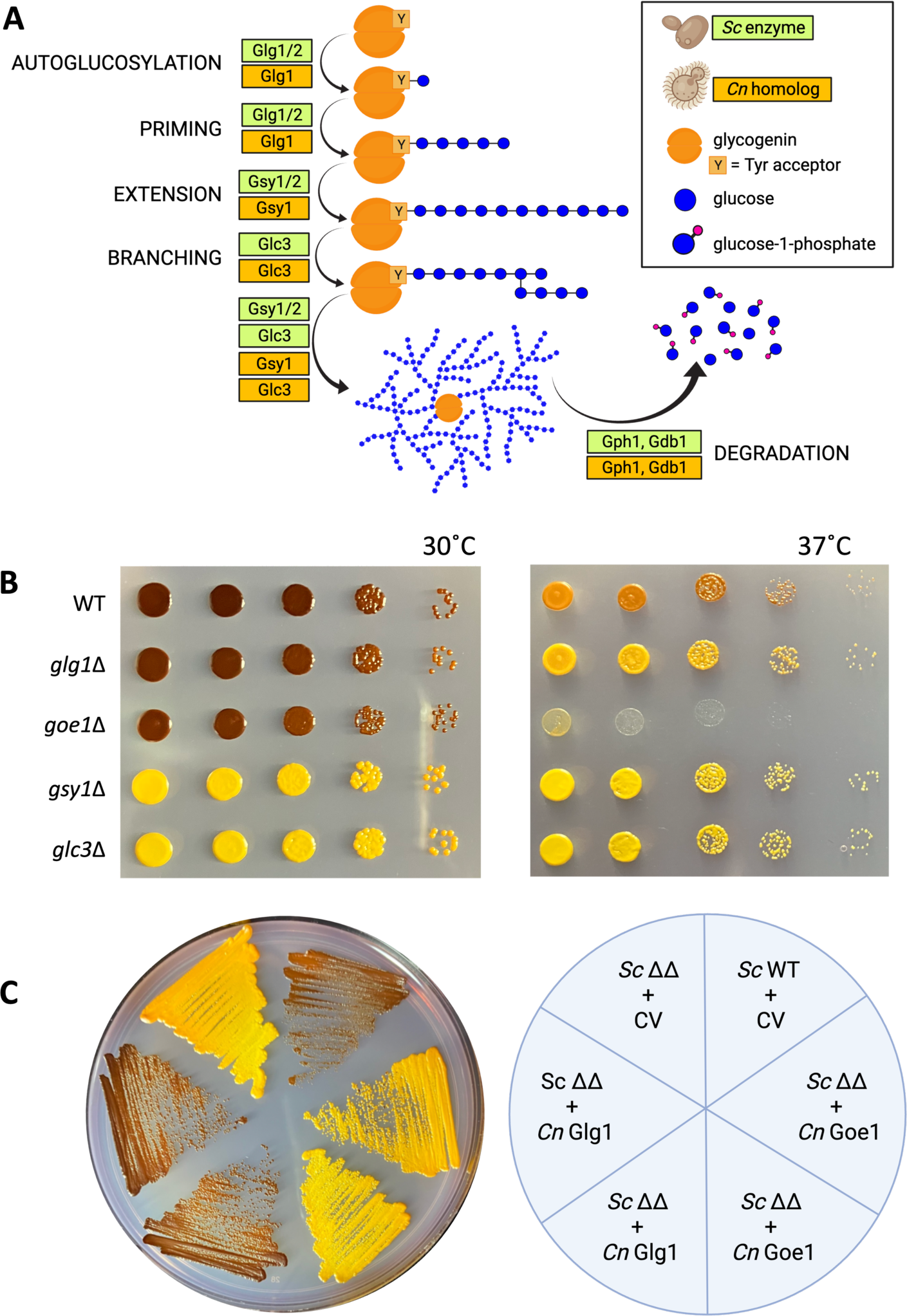
Glycogen synthesis in *S. cerevisiae* (*Sc*) and *C. neoformans* (*Cn*) A. Schematic of glycogen synthesis and degradation in *Sc* and *Cn*. B. Serially diluted *Cn* cells of the indicated strains, grown on SD+Gal for 72 h and exposed to iodine vapor. C. *Sc glg1 glg2* (*Sc* ΔΔ) or WT cells were transformed as indicated at right with control vector (CV) or plasmids encoding *Cn* Glg1 or Goe1 (two independent transformants for each). Transformants were grown on SD-Trp+Gal for 72 h and exposed to iodine.

Carbon levels drive the expression, phosphorylation, and allosteric regulation of enzymes responsible for glycogenesis and glycogenolysis. Tight coordination of these regulatory processes allows cells to adapt to the vagaries of carbon availability in their immediate environment. If extracellular carbon is low, glycogen is degraded by the activities of glycogen phosphorylase (Gph1) and debranching enzyme (Gdb1) (Figure 1A). The resulting Glc-1-phosphate (Glc-1-P) and free Glc can then be used elsewhere in the cell^10–12^.

For pathogens that must inhabit multiple niches with varying nutrient availability, both outside and within a host, carbohydrate storage is critical. The focus of this work, *Cryptococcus neoformans* (*Cn*), is an opportunistic fungal pathogen that primarily afflicts those living with HIV/AIDS, killing more than 112,000 people globally each year^13^. *Cn* encounters diverse environments during its life cycle, ranging from soil, trees, and bird excreta to infected mammals^14–16^. When infectious particles are inhaled into the lungs of a suscepti-ble host, the fungi survive phagocytosis by innate immune cells^17^. Free and intracellular cryptococci may then disseminate via the bloodstream and cross the blood-brain barrier to cause an often-fatal meningitis^18–20^.

At each site of infection within the host, fungi encounter a unique set of environmental conditions. Notably, glucose is abundant in the bloodstream and central nervous system but limited in the lung and inside phagocytes, highlighting the need for reserve carbohydrates^21–24^. *Cn* makes two intracellular storage poly-mers: trehalose and glycogen. Trehalose (an α-1,1-linked Glc disaccharide) is important for sporulation and pathogenesis^25–27^. Glycogen has not been studied in Cn, other than one report that it is dispensable in the chlamydospore^28^.

In addition to glucose storage molecules, *Cn* incorporates a variety of glucose polymers into its cell wall. In fungi, the cell wall is a dynamic barrier that helps the cell resist external stressors such as UV light (in the environment), reactive oxygen species (in a host), and changes in temperature, pH, and osmolarity. The cryptococcal cell wall features a dense inner layer of chitin, made by a family of synthases and modified by a family of deacetylases, and long β-1,3-glucan polymers synthesized by Fks1^29–32^. Although *FKS1* is essential, cells with reduced expression remain viable by increasing cell wall chitin^31,33^. Such compensatory mechanisms may also explain intrinsic cryptococcal resistance to echinocandin drugs, which effectively target Fks1 in other fungi^34,35^. The less dense outer layer of the wall contains primarily α-1,3-glucan, which is assembled by _Ags136,37._

In both wall layers, abundant short β-1,6-glucans link the other glycans to each other and to cell wall proteins^38^. The synthesis of β-1,6-glucan is poorly understood. In *Sc*, this process has been localized to the plasma membrane^39^. However, proteins in the ER and Golgi (Kre5, Kre6, and Skn1^40–42^), cytosol (Kre11^43,44^), and at the cell surface (Kre1, Kre9, and Knh1^45–47)^ all affect its abundance, suggesting that β-1,6-glucan syn-thesis involves both intracellular and extracellular steps. Gas1 and Bgl2 activities, respectively located in the plasma membrane and cell wall, introduce β-1,6 linkages in branched β-1,3-glucan^48–50^. Cryptococcal versions of the ER/Golgi proteins Kre5, Kre6, and Skn1 have been characterized^51^. However, no cell surface proteins have been implicated in β-1,6-glucan synthesis in this organism.

Also present in the cell wall is a second form of α-1,4-glucan, distinct from intracellular glycogen in its localization and physical properties. Termed ‘insoluble glycogen’ in a small body of literature dating back to 1925^52^, this material has been best studied in *Sc*^53–57^ and recently reported in *Candida* species^58^. Detailed structural analyses have shown that cell wall α-1,4-glucan is covalently bound to β-1,3-glucan via a β-1,6-glucan linker^56,58^, mirroring the connections of other wall components. Unlike intracellular glycogen, it con-tains no protein component^58^. Though never directly studied in *Cn*, α-1,4-glucan has been observed inci-dentally as a minor component (1%) of the cryptococcal cell wall^59^. To date, no enzymes involved in the syn-thesis or trafficking of cell wall α-1,4-glucan have been characterized, in any system.

We characterized a glycogenin that is responsible for initiating glycogen synthesis in *Cn*. In the absence of this protein, no α-1,4-glucan is detected in the cell wall, which suggests that cell wall α-1,4-glucan is de-rived in some way from intracellular glycogen. In our search for related proteins, we also discovered a novel glycosyltransferase involved in both glycogen and cell wall synthesis. This enzyme affects the abundance and solubility of β-1,3-glucan in the cell wall. Altogether, our studies have revealed unexpected links between glycogen and cell wall synthesis, with important implications for fungal pathogenesis.

## RESULTS

### Glycogen synthesis in Cn

Inspired by the idea that *C. neoformans* (*Cn*) might utilize glycogen strategically as part of its patho-genic lifestyle, we investigated cryptococcal genes predicted to encode the glycogen synthetic pathway. *S. cerevisiae* (*Sc*) employs two glycogenins and two glycogen synthases. In *Cn*, by contrast, only one glycogenin (Glg1, encoded in the KN99α wild-type strain by *CKF44_05293*) and one synthase (Gsy1, encoded by *CKF44_04621*) are predicted by homology to their yeast counterparts (33% and 64%, respectively). Like *Sc*, *Cn* has a single branching enzyme (Glc3, encoded by *CKF44_00393*). Looking for related proteins within the *Cn* genome, we found that *CKF44_02199* encodes a hypothetical glycosyltransferase that is 35% identical at the amino acid level to *Cn* Glg1 (it has no similarity to *Sc* Glg1 or Glg2). We named this gene <u>G</u>lucan <u>o</u>rganizing <u>e</u>nzyme 1 (*GOE1*), for reasons that will be discussed below.

To investigate glycogen synthesis in *Cn*, we engineered or obtained mutants lacking synthetic genes (see Methods and Acknowledgements). We grew these strains on synthetic defined medium with galactose as the sole carbon source (SD+Gal) and exposed them to iodine vapor, which stains intracellular glycogen brown^28^ (Figure 1B). The absence of glucose in this medium induces glycogenolysis, allowing us to see varia-tions in glycogen abundance between strains. In this assay, *gsy1*Δ and *glc3*Δ, which are impaired in late stages of synthesis (Figure 1A), were strikingly deficient in glycogen at both temperatures tested; colonies appeared bright yellow. Interestingly, at 37°C, *glg1*Δ was slightly darker than *gsy1*Δ and *glc3*Δ, despite the presumed role of Glg1 as the sole initiator of glycogen synthesis. *goe1*Δ also exhibited a glycogen defect at 37°C, as well as a severe growth defect. In contrast, both *glg1*Δ and *goe1*Δ contained abundant glycogen at 30°C, implying the involvement of as-yet-unidentified proteins in glycogen synthesis at the lower temperature. However, we limited our studies to physiological temperature due to our focus on cryptococcal pathogenesis.

To test whether Glg1 and Goe1 were functional glycogenins, we expressed each protein in a *Sc* strain that lacks both glycogenins (*Sc glg1 glg2*; Figure 1C). Cryptococcal Glg1 completely restored glycogen synthe-sis in *Sc glg1 glg2*, confirming that it is indeed capable of autoglucosylation and initiation of glycogen synthe-sis. In contrast, expression of *GOE1* did not rescue glycogen synthesis in this system; Goe1 is therefore unlikely to be a second cryptococcal glycogenin.

### Perturbations of *GLG1* impact glycogen levels

To further characterize Glg1 in *Cn*, we created point mutations in two key conserved sites^60^. The first mutation, K094A (Figure 2A, red arrow), changes a residue that is thought to interact with a phosphate group of the nucleotide sugar donor UDP-Glc^61^. Mutation of this site in rabbit glycogenin abolishes catalytic activ-ity^61^, so we refer to it as the catalytic site. Complementation of *glg1*Δ cells with the catalytic mutant version of the protein in the endogenous locus (*glg1*Δ::*GLG1^cat^*) phenocopied the deletion mutant in our iodine assay (Figure 2B, compare second and fourth rows), while similar complementation with the unperturbed sequence phenocopied WT (compare first and third rows). To measure glycogen content, we used an assay based on the specificity of amyloglucosidase for terminal α-1,4-Glc linkages. Briefly, the supernatant fraction (16,000 rcf) of lysed cells was treated with this enzyme and any released glucose detected by a colorimetric probe. As in the iodine assay, we observed less glycogen in *glg1*Δ and *glg1*Δ::*GLG1^cat^* than in WT and complemented cells (Figure 2C).

**Figure 2.**
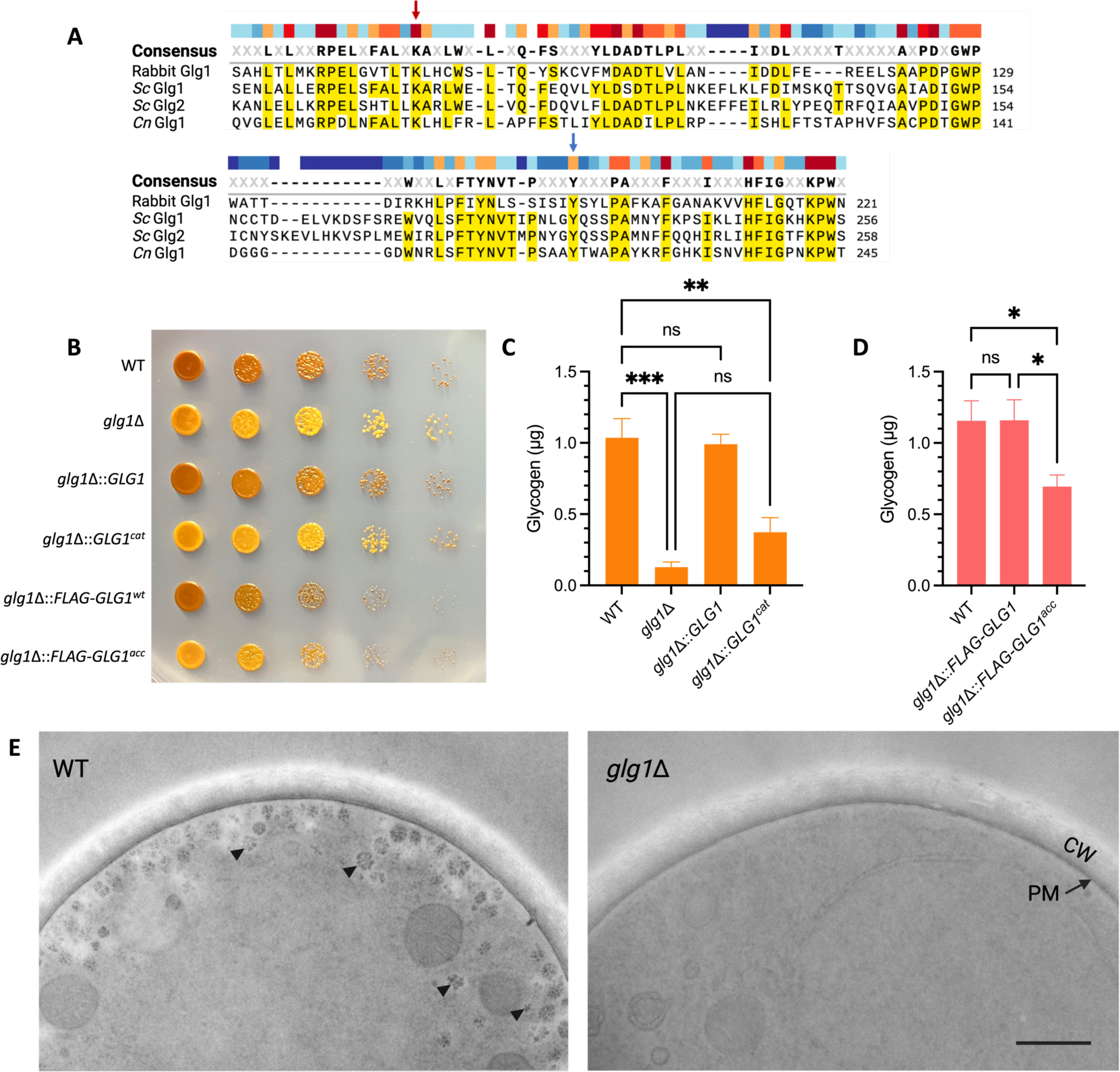
Mutation of Glg1 impacts glycogen levels. A. Partial T-Coffee alignments of glycogenin and Goe1 sequences^60^. Red arrow, conserved lysine; blue arrow, acceptor tyrosine. B. Iodine assay of the indicated strains as in Fig. 1. C-D. Glycogen content by enzymatic assay. Data are shown as mean + SEM of 3 (C) or 6 (D) biological replicates; ns, p>0.05; *, p<0.05; **, p<0.01; ***, p<0.001 by ANOVA. Limits of detection (LOD), 0.043 µg (C) and 0.038 µg (D). E. Electron micrographs of Thiery-stained cells. CW, cell wall; PM, plasma membrane; arrowheads, example rosettes. Images are to the same scale; scale bar, 500 nm.

The second mutation, Y219A (Figure 2A, blue arrow), corresponds to the residue that accepts the initial glucose during autoglucosylation in the *Sc* glycogenins (acceptor site). In the acceptor mutant strain, we also incorporated an N-terminal 3xFLAG tag (*glg1*Δ::*FLAG-GLG1^acc^*). *glg1*Δ::*FLAG-GLG1^acc^* clearly had re-duced glycogen compared to its control strain *glg1*Δ::*FLAG-GLG1^wt^* by iodine assay (Figure 2B). This observa-tion was borne out in enzymatic assays (Figure 2D).

Glycogen granules assemble into rosettes in several cell types, including yeast^62,63^. To visualize glyco-gen in WT and *glg1*Δ cells, we subjected them to freeze-substitution electron microscopy and Thiery staining as in Coulary *et al*^63^. In WT cells, glycogen rosettes were clearly visible and were clustered at the periphery of the cytosol near the plasma membrane (Figure 2E; see Figure S1 for more images). By contrast, *glg1*Δ cells were completely devoid of rosettes but otherwise appeared normal.

### Deletion of *GOE1* impacts glycogen abundance, rosette localization, and cell wall integrity

Having established Glg1 as a canonical glycogenin of particular importance at 37°C, we turned our attention to the novel protein Goe1. As noted above, this protein did not complement glycogenin deficiency in *Sc* (Figure 1C), yet it seemed to have a role in both glycogen synthesis and cell growth (Figures 1B and 3A). The glycogen defects observed in *glg1*Δ and *goe1*Δ (Figure 1B) were exacerbated when both genes were deleted (Figure 3A), hinting that these proteins might operate in the same pathway. Complementation of *goe1*Δ with WT *GOE1* (*goe1*Δ::*GOE1*) restored glycogen levels and normal growth.

**Figure 3.**
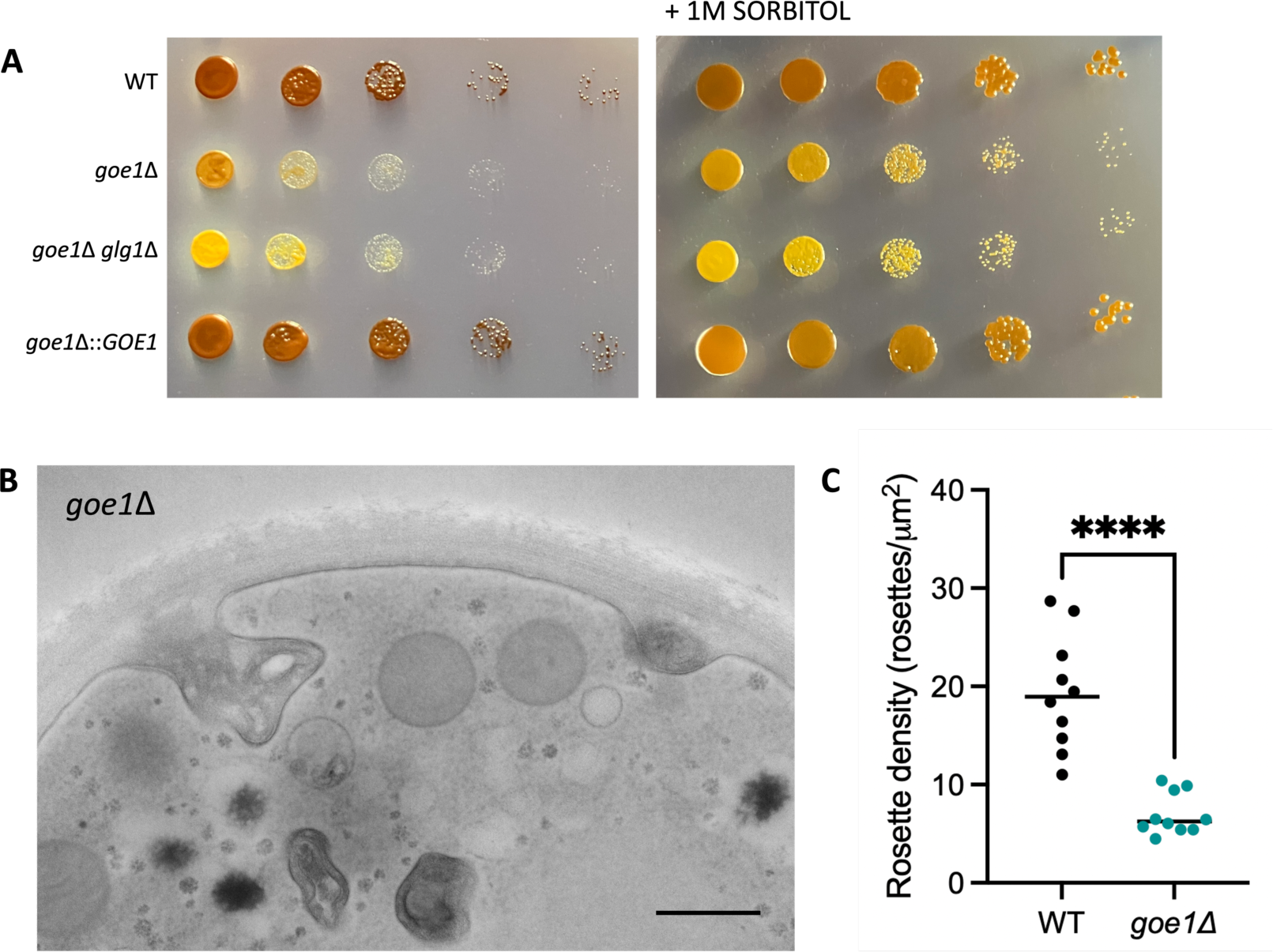
Goe1 affects glycogen abundance, roseRe localization, and cell wall integrity. A. Iodine assay of the indicated strains grown on SD+Gal α sorbitol for 96 h. B. Electron micrographs of Thierystained cells. Scale bar, 500 nm. C. QuanWtaWon of rose]e density within 500 nm of the cytosol-PM interface (see Methods). Symbols, value for individual images (10 per strain); black bar, mean; ****, p<0.0001 by t-test.

EM coupled with Thiery staining yielded more surprises in the *goe1*Δ deletion mutant. In striking con-trast to WT images, the glycogen rosettes in these cells were scattered throughout the cytosol (compare Figure 3B to 2E), such that their density close to the plasma membrane was reduced to 36% of WT (Figure 3C and Figure S2). This mutant also showed dramatic cell wall disorganization and invaginations that were not present in WT and *glg1*Δ cells (Figure 3B and Figure S2). Overall, these images implicate Goe1 in both locali-zation of glycogen rosettes and cell wall integrity.

### Topology and localization of Goe1

All known glycogen synthetic enzymes are soluble. In this context, it was puzzling to us that Goe1, which clearly affects glycogen synthesis, is predicted to contain a transmembrane domain. Moreover, the temperature sensitivity, largely rescued by sorbitol (Figure 3A), and electron micrographs of *goe1*Δ cells suggest that Goe1 is involved in cell wall construction. To reconcile these findings, we hypothesized that Goe1 resides in the plasma membrane, where it interacts with glycogen and/or its associated enzymes as well as with the cell wall.

To investigate the localization and orientation of Goe1, we first attempted to localize a version of Goe1 tagged with mNeonGreen, both directly and via immunofluorescence, but were unable to do so. We next turned to another strategy, biotin labeling. Sulfo-NHS-LC-LC-biotin is a membrane-impermeant compound that irreversibly labels accessible lysine residues or N-termini of proteins, increasing their molecular weight by 452.6 Daltons per biotin added. When it is incubated with intact cells, only the extracellular domains of proteins are biotinylated. This approach exploits the asymmetry of Goe1: its N-terminus consists of 19 amino acids including 2 lysines before the transmembrane domain, while the C-terminal remainder of the protein (321 amino acids) features 24 lysines (Figure 4A). If the N-terminus of Goe1 is extracellular, the molecular weight of Goe1 would increase by a maximum of ∼1.4 kD upon labeling. If the C-terminus is extracellular, the protein could increase by up to ∼10.9 kD.

**Figure 4.**
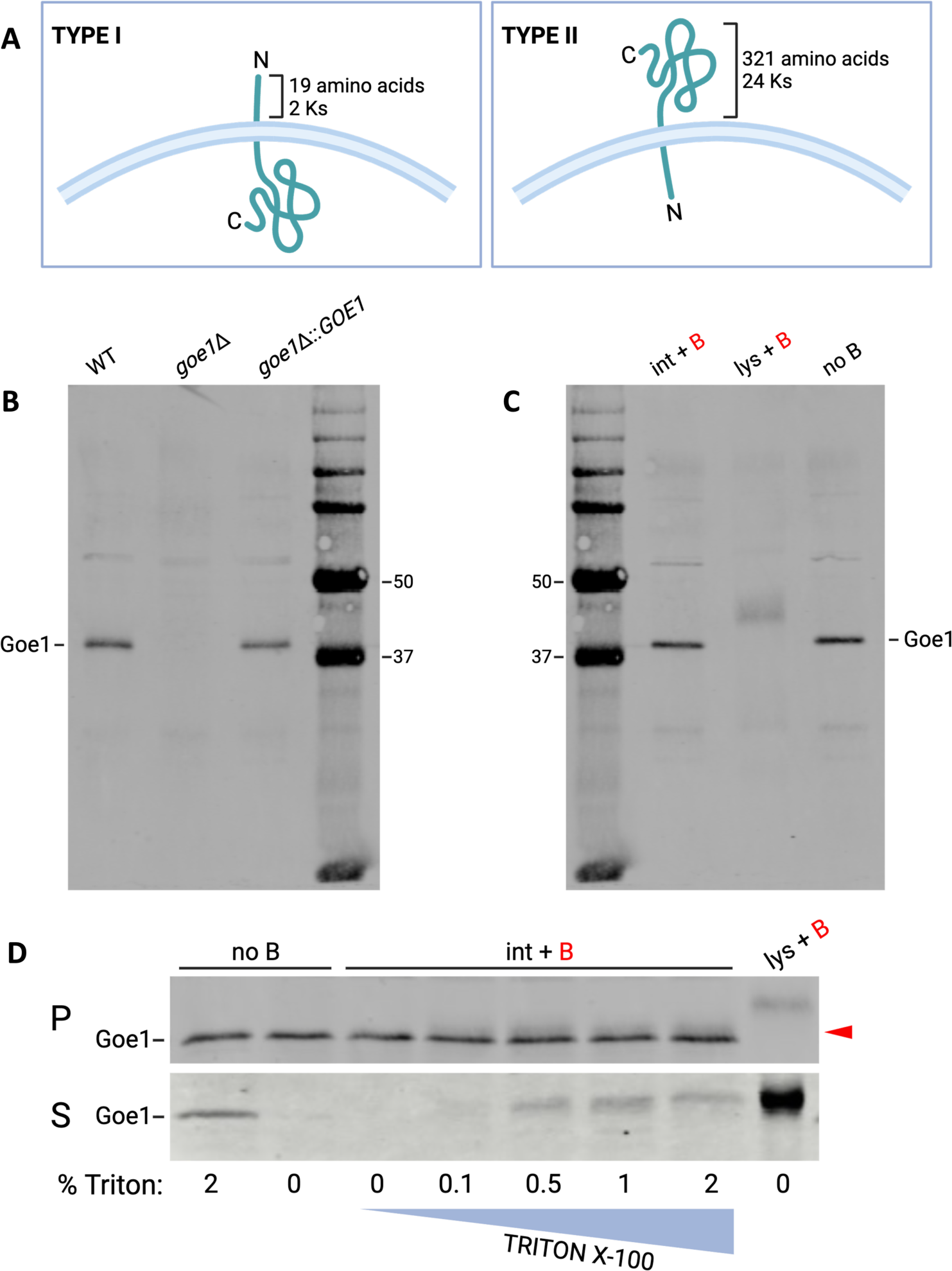
Goe1 is likely a plasma membrane protein with an extracellular N-terminus. A. Sche-matic of possible Goe1 topologies. B. α-Goe1 immunoblot of indicated strain lysates, 25 µg total protein loaded per lane. C. α-Goe1 immunoblot of lysates (25 µg per lane) from WT cells either incubated with biotin (int + B), lysed prior to incubation with biotin (lys + B) or never exposed to biotin (no B). D. α-Goe1 immunoblot of samples labeled as above. Cells were treated with Triton X-100 at the indicated % prior to biotin labeling. P, pellet fractions (25 µg/lane); red triangle, bioti-nylated species of Goe1 (see text). S, supernatant fractions (16 µL/lane), except final lane (25 µg pellet for size reference).

A polyclonal rabbit antibody raised against amino acids 91-366 of Goe1 (see Methods) revealed the protein at a migration position consistent with its predicted molecular weight of 42 kD in both WT and *goe1*Δ::*GOE1*. This band is absent in *goe1*Δ, demonstrating antibody specificity (Figure 4B). When intact cells were labeled with biotin, the migration of the Goe1 band was similar to that of native Goe1 (Figure 4C, com-pare first and last lanes). However, when cells were lysed before labeling, we observed a range of mobility that was higher on the gel than native Goe1 (Figure 4C, middle lane). This result suggested that the large catalytic C-terminus of Goe1 may be intracellular.

It was unclear whether biotin labeling of Goe1 occurred in intact cells, since we did not observe a shift in migration compared to non-biotinylated Goe1. One possibility was that the N-terminus was labeled, but that this was difficult to discern by immunoblot due to the small change in mass. Alternatively, the labeling was unsuccessful due to inaccessibility of the extracellular domain, which may be in complex with other pro-teins. To address the latter possibility, we incubated intact cells with increasing amounts of Triton X-100 (TX-100) prior to biotin labeling. TX-100 is a nonionic surfactant that, depending on concentration, may disrupt protein complexes *in situ* and extract proteins from the plasma membrane^64,65^. After detergent treatment and labeling, we subjected the samples to centrifugation, reserved the supernatant fractions, and lysed the pellet fractions.

In pellet fractions (Figure 4D, top panel) of cells that had been incubated with TX-100 prior to biotin labeling (marked 0.1-2%), we reproducibly observed a faint Goe1 species ∼1 kD larger than native Goe1 (red triangle). This new band appeared consistently across the range of detergent treatment but was absent from cells treated with 2% TX-100 alone (first lane) or cells labeled without Triton (third lane). We interpret this band to be plasma membrane-associated Goe1 with a biotinylated and hence extracellular N-terminus, made accessible by TX-100 treatment.

In supernatant fractions (Figure 4D, bottom panel), we observed the emergence of a higher molecular weight species of Goe1 in samples treated with TX-100. This band was similar to that seen when cells were lysed before incubation with biotin (right-most lane), suggesting that some Goe1 was fully extracted from the membrane by TX-100 and maximally biotinylated in this experiment.

### Cell wall α-1,4-glucan

Given our evidence for its dual involvement with glycogen and the cell wall, we speculated that Goe1 either participated in the synthesis of α-1,4-glucan destined for the cell wall, or influenced its incorporation into the cell wall. To pursue this idea, we used methods from the prior literature on ‘insoluble glycogen’^57^ to isolate the alkali-insoluble cell wall fraction and then measured its α-1,4-glucan content enzymatically as above. This fraction is traditionally understood to contain chitin and any glycans covalently linked to it, either directly or indirectly (e.g. β-1,3-glucan connected to chitin via β-1,6-glucan segments).

These studies yielded the exciting observation that the walls of WT cells indeed contained α-1,4-glu-can (Figure 5A), confirming a single early report in the literature^59^. This material was undetectable by enzy-matic assay in *glg1*Δ cells, which indicates that α-1,4-glucan in the cell wall is derived from intracellular gly-cogen and not via a separate synthetic pathway. Compared to WT, it was reduced 8-fold in *goe1*Δ and was restored in the complemented mutant. Notably, while both *goe1*Δ and *glg1*Δ cells were deficient in cell wall α-1,4-glucan in this assay, only *goe1*Δ exhibited aberrant cell wall morphology.

**Figure 5.**
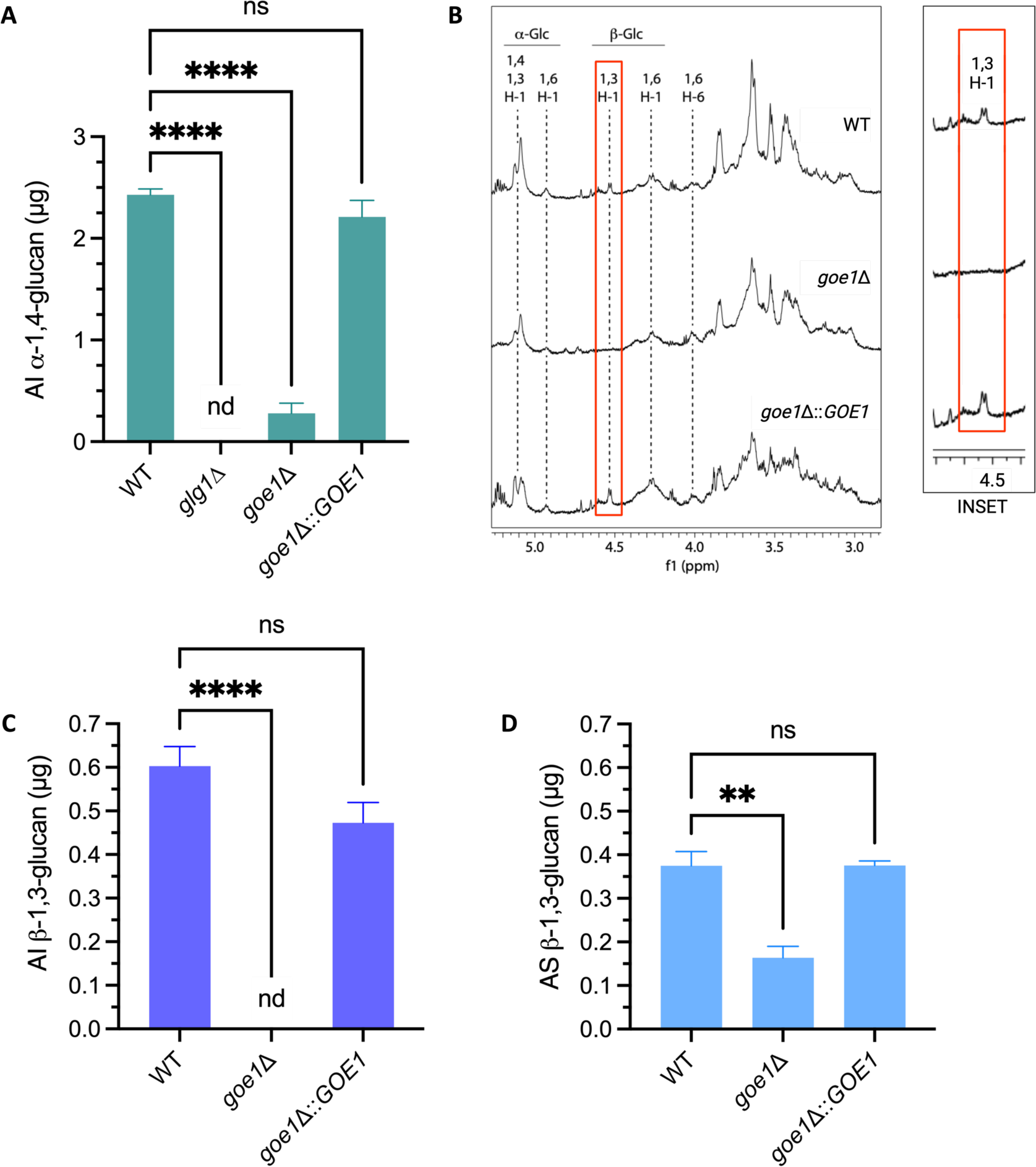
Goe1 affectsα-1,4 and β-1,3-glucans in the cell wall. A. Enzymatic quantitation of α-1,4-glucan in the alkali-insoluble cell wall fraction. Data are shown as mean + SEM of three biological replicates; ***, p<0.001; ****, p<0.0001 by ANOVA; LOD = 0.058. B. NMR spectra of alkali-insoluble cell wall fractions. Red box and inset, β-1,3-glucan peak. The signals in the region between 3 and 4 ppm on the right side of the spectra correspond to the carbohydrate ring protons and cannot be assigned based on 1D proton NMR alone. The peaks near 5.4 ppm are of unknown origin. C-D. Enzymatic quantitation of β-1,3-glucan in the alkali-insoluble (AI, Panel C) or alkali-soluble (AS, Panel D) fractions of the indicated strains. Mean + SEM of three biological replicates shown; **, p<0.01 by ANOVA; LOD = 0.046 in D; LOD = 0.081 in E.

To explore the cell wall defects of *goe1*Δ, we analyzed the alkali-insoluble fraction using 1D NMR. Unfortunately, this did not help us understand α-1,4-glucan because this NMR method did not allow specific peak assignment for this linkage (Figure 5B, signals near 5.2 ppm). Linkage analysis performed in parallel suggested an increase in 4-linked glucan (Table S1), in contrast to our enzymatic studies (Figure 5A); this dis-crepancy may relate to altered susceptibility of α-1,4-glucan to amyloglucosidase digestion in *goe1*Δ cells.

### Cells without *GOE1* lack β-1,3-glucan in the alkali-insoluble cell wall fraction

To our surprise, the NMR studies revealed a striking loss of the anomeric signal corresponding to β-1,3-glucan in *goe1*Δ cells (Figure 5B, red box and inset). Consistent with this finding, 3-linked and 3,6-linked glucopyranosyl residues were reduced in this fraction of *goe1*Δ cells compared to WT and *goe1*Δ::*GOE1* (Table S1). To test this by a third independent method, we used an enzymatic assay; again, no β-1,3-glucan was detected in the alkali-insoluble fraction of *goe1*Δ (Figure 5C). Interestingly, some β-1,3-glucan was present in the alkali-soluble fraction (containing glycans not linked to chitin) of *goe1*Δ, though less than in WT (Figure 5D; see Discussion). In both fractions, we detected normal levels of β-1,3-glucan in the complemented strain *goe1*Δ::*GOE1*.

### Glg1 and Goe1 are implicated in pathogenesis

Cryptococcal pathogenesis hinges on the interactions between fungi and alveolar macrophages in the earliest stages of infection. We speculated that glycogen, in its traditional role of energy reserve, could coun-ter the phagocytic strategy of carbon starvation^66^. Goe1 also contributes to cell wall integrity, a bulwark against reactive oxygen species produced by host cells. We therefore evaluated fungal survival in THP-1 mac-rophage-like cells for both *glg1*Δ and *goe1*Δ cells. Both strains were phagocytosed normally, with 1,400-3,200 cryptococci engulfed per well (Figure 6A, 0-hour timepoint). To our surprise, *glg1*Δ cells proliferated as well as WT after engulfment, indicating that glycogen is not required in this particular niche (Figure 6A). However, *goe1*Δ cells exhibited a notable intracellular survival defect.

To further assess our mutants in the more complex environment of the lung, we measured fungal burden 9 days after intranasal inoculation of mice. We found that infection with *glg1*Δ yielded a roughly 4-fold reduction in total lung burden, which was restored in the complemented mutant (Figure 6B). Infection with *goe1*Δ also resulted in significantly lower lung burden compared to controls, with a reduction of 28-fold compared to WT for the single mutant. The double mutant remained close to the level of inoculation, a 59-fold decrease in lung burden compared to WT (Figure 6C).

**Figure 6.**
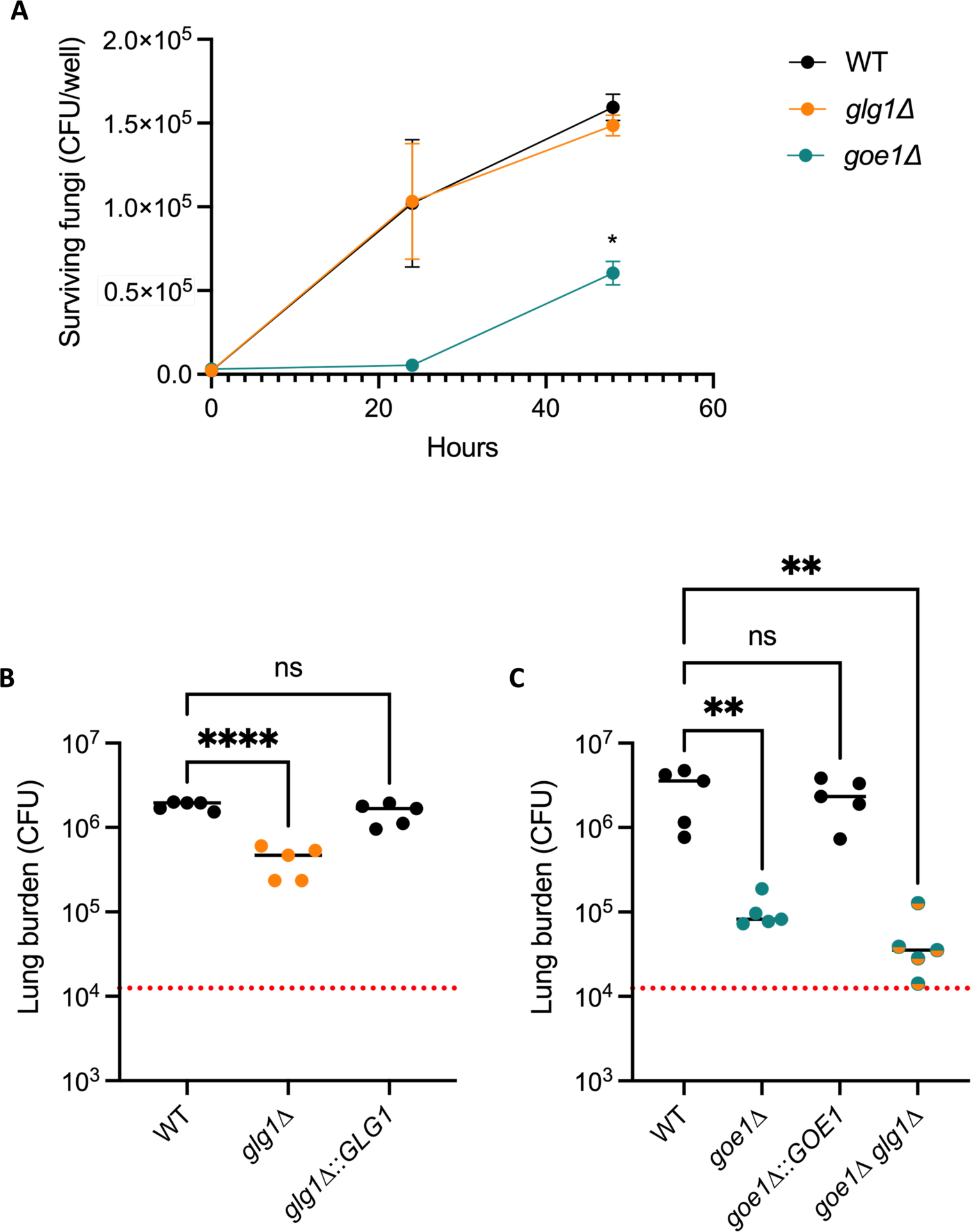
Loss of Glg1 or Goe1 reduces cryptococcal pathogenesis. A. Intracellular survival in THP-1 cells shown as mean α SEM of two biological replicates; *, p<0.05 by two-way ANOVA; CFU, colony forming units. B-C. Lung burden resulting from 9-day infection with 1.25×10^4^ cells of each indicated strain. Data are shown as mean α SEM shown; **, p <0.01; ****, p <0.0001 by ANOVA. Red do]ed line indicates initial inoculum.

## DISCUSSION

Our initial interest in glycogen led us to intriguing discoveries about the *C. neoformans* cell wall, which raise multiple questions for future research. We first established that Glg1 is a canonical glycogenin, capable of initiating glycogen synthesis, and appears to be the only one encoded by the cryptococcal genome. However, in its absence, a limited amount of glycogen is still made at 37°C, and glycogen is abundant at 30°C.

We therefore posit the existence of another substrate upon which glycogen can be built, an idea first suggested in regard to *Sc*^67^. We hypothesize that glycogen synthase and branching enzyme can build branched chains on this unknown acceptor, generating an alternate product that is stained by iodine in *glg1*Δ but is not present in *gsy1*Δ or *glc3*Δ cells. Interestingly, this product does not form rosettes in our growth conditions, as seen in our EM images of *glg1*Δ. Further studies will elucidate the details of this alternate synthesis pathway.

Beyond its role as the major component of glycogen, α-1,4-glucan is also an often-overlooked component of the fungal cell wall. One previously untested assumption is that this material is derived from intra-cellular glycogen^58^. The absence of detectable cell wall α-1,4-glucan in *glg1*Δ cells strongly supports the idea that this material indeed originates via the glycogen synthesis pathway.

In our search for novel enzymes involved in glycogen synthesis, we discovered Goe1, which is the cryptococcal protein most similar to Glg1. Unlike Glg1, however, Goe1 is membrane bound. Our biotin labeling studies suggest that Goe1 is a type 1 plasma membrane protein, with its C-terminal domain in the cytosol. However, our results do not definitively rule out other interpretations. For example, the C-terminus could be extracellular but poorly accessible to biotinylation, even in the presence of detergent.

Goe1 has at least two distinct biological roles. First, it is responsible for the localization of glycogen rosettes to the plasma membrane (compare Figures 2E and 3B); this tethering could be mediated by Goe1 interacting directly with glycogen, with another protein, or both. This association may increase the efficiency of glycogen synthesis by bringing rosettes (and their accompanying enzymes) together spatially, explaining the glycogen defect of *goe1*Δ cells. Plausibly, fungal glycogen serves not only as an energy reserve, but also as a source of building materials for the cell wall, since its primary breakdown product (Glc-1-P) may be used to form the donor for glucan synthesis (UDP-Glc). Proximity of these reactions to synthases in the plasma membrane could facilitate efficient synthesis of cell wall glucans.

The second role of Goe1 is related to its putative glycosyltransferase activity. The phenotypes of *goe1*Δ cells strongly implicate its enzymatic function in cell wall synthesis or remodeling. The inner layer of the cell wall is particularly disrupted by the loss of Goe1 (e.g. Figure 3B and Figure S2), hinting that its product might be found there. This layer is composed of chitin/chitosan along with α-1,4-glucan, β-1,6-glucan, and β-1,3-glucan. We tested one hypothesis, that Goe1 incorporates α-1,4-glucan into the cell wall, by measuring α-1,4-glucan in the cell walls of our mutants. While both *goe1*Δ and *glg1*Δ cells had reduced cell wall α-1,4-glucan, only *goe1*Δ exhibited dramatic cell wall disorganization. This defect must therefore be caused by the loss of a different cell wall component.

1D proton NMR of the alkali-insoluble fractions of our strains provided important clues. Levels of β-1,6 glucan in *goe1*Δ appeared normal in our NMR and linkage analyses. Goe1 is therefore unlikely to be the main enzyme that generates β-1,6-glucan. In contrast, the spectrum for *goe1*Δ completely lacked signal from β-1,3-glucan, a finding we confirmed by enzymatic assay. The latter did detect β-1,3-glucan in the alkali-soluble fractions from this mutant, though at lower levels than in WT. Taken together, these findings lead to our working hypothesis, that Goe1 connects β-1,3-glucan and β-1,6-glucan in the cell wall. In the absence of Goe1, β-1,3-glucan is still synthesized and extruded by Fks1, but without its linkage via β-1,6-glucan to other cell wall components, it ends up in the soluble fraction. As a result, the wall structure is perturbed, and cell wall integrity becomes compromised.

Evidence in other fungi suggests that the addition of mature β-1,6-glucan polymers to β-1,3-glucan occurs extracellularly, following β-1,3-glucan branching mediated by the GPI-anchored Gas/Gel/Phr family^50,68^. This contradicts our proposed topology for Goe1. However, in contrast to other fungi, the dominant β-linked cell wall glucan of *Cn* is β-1,6 rather than β-1,3^59^. Therefore, synthesis of its wall might require unique cross-linking pathways.

We can imagine at least two possible enzymatic activities for Goe1 that are consistent with our data. First, Goe1 could add one or more glucose moieties in β-1,6-linkage upon a β-1,3-glucan scaffold, a process that could conceivably occur on either side of the plasma membrane, depending on protein topology. This nascent β-1,6 branch could later serve as an acceptor for other β-1,6-glucan synthase or glucanosyltransferase enzymes. By this model, Goe1 makes the linkage required for β-1,3-glucan to become insoluble. Alterna-tively, Goe1 could itself act as a glucanosyltransferase, adding an already-synthesized β-1,6-glucan branch to β-1,3-glucan or vice versa. This would be most likely if the protein C-terminus is actually outside of the cell. Differentiating between these models will require direct demonstration of Goe1’s enzymatic activity.

The present study begins to illuminate unexplored pathways of cell wall synthesis related to glycogen, generating exciting new questions for the field. We have confirmed the presence of α-1,4-glucan in the cryptococcal cell wall, likely derived from intracellular glycogen, but the mechanism of this transfer remains to be defined. Future work on Goe1 will be needed to determine how it organizes glycogen rosettes and whether its glycosyltransferase activity indeed links β-1,3-glucan and β-1,6-glucan.

## MATERIALS AND METHODS

### Strain construction & cell growth

To generate mutant and complemented mutant strains in *Cn,* we utilized a split-marker strategy with biolistic transformation^69^. First, we replaced the complete coding sequence of *GLG1* with a Geneticin (G418) resistance cassette by biolistic transformation of the KN99α strain to create *glg1*Δ. We complemented this strain at the native locus with WT or mutated gene sequences using the same method, replacing the G418 marker with a cassette containing *GLG1* and a nourseothricin (NAT) resistance marker in tandem. We used the same approach to modify the *GOE1* locus in KN99**a**, using NAT and G418 to mark the deletion and complemented strains, respectively. To obtain a *glg1*Δ *goe1*Δ double mutant, we crossed the single mutants on V8 agar plates^70^, selected strains resistant to both drugs, tested mating type using colony PCR^71^, and used a MAT**a** strain in our studies. All strain manipulations were checked by acquired drug resistance and PCR, then con-firmed by whole-genome sequencing^72^.

For all studies, *Cn* and *Sc* strains were inoculated from single colonies into the medium indicated. Cells were grown at the temperature indicated in either tubes (5 mL cultures) or baffled flasks (25 mL cultures) with shaking at 230 rpm.

### Heterologous expression in *S. cerevisiae*

The coding sequences of *GLG1* and *GOE1* were each amplified from cDNA and cloned into the yeast expres-sion vector pD1215 using the Electra system (Atum). We also generated a control vector containing the *GAL4* coding sequence. A yeast strain lacking both glycogenin genes (CC9; here *Sc glg1 glg2*) and its isogenic wild-type control (EG328-1A; *Sc* WT) were generous gifts from Wayne Wilson at Des Moines University^4^. *Sc* strains were transformed using the lithium acetate method^73^ and plated on tryptophan dropout medium (SD-Trp+Glc: 0.66% w/v YNB without amino acids, 0.08% w/v amino acid mix lacking tryptophan, 2% w/v agar, 2% w/v glucose).

### Iodine assay

Cryptococcal strains were grown overnight in 5 mL YPD (1% w/v BactoYeast Extract, 2% w/v BactoPeptone, 2% w/v dextrose) and then washed and adjusted to 10^7^ cells/ml in sterile PBS. This stock was first diluted 10-fold to a concentration of 10^6^ cells/ml, which was then serially diluted 5-fold three more times. Cells at each of these five concentrations were spotted on synthetic defined media with galactose (SD+Gal: 0.66% w/v YNB without amino acids, 2% w/v galactose, 2% w/v agar). *Sc* transformants were streaked onto tryptophan drop-out medium lacking Glc (SD-Trp+Gal: 0.66% w/v YNB without amino acids, 0.08% w/v amino acid mix lacking tryptophan, 2% w/v agar, 2% w/v galactose). All plates were incubated for 72-96 h to allow fungal growth. Unless otherwise stated, *Cn* cells were grown at 37°C while *Sc* cells were grown at 30°C. Lidless plates were then transferred to a covered glass dish containing evenly distributed iodine crystals (2 g) and photographed after 30 min of iodine exposure.

### Glycogen assay

Strains were grown for 6 h in 5 mL YPD at 37°C, then subcultured into 25 mL SD+Gal and grown for a further 18 h at 37°C. Samples of 10^8^ cells were resuspended in 500 µL water, mixed with 300 µL silica beads, and subjected to two 90-sec rounds of bead-beating, incubating for 1 min on ice between rounds. Lysates were collected and centrifuged (16,000 rcf; 30 min; 4°C); the resulting supernatant fractions were boiled for 10 min to inactivate enzymatic activity and their glycogen content measured using a Glycogen Assay Kit (ab65620, Abcam). Briefly, in this assay glycogen in samples and standards is subjected to hydrolysis by amylogluco-sidase, which specifically cleaves α-1,4 glycosidic bonds from the nonreducing end of an α-1,4 glucan chain. The resulting free glucose is subsequently oxidized so that it can react with a colorimetric probe. Each reaction contained 5 μg of protein and all samples and standards were assayed in triplicate. Absorbance (570 nm) was measured using a BioTek plate reader. To control for background cellular glucose levels, the average absorb-ance of sample reactions lacking glucoamylase was subtracted from that of the corresponding sample reac-tions containing amyloglucosidase. Glycogen content of background-corrected samples was calculated by in-terpolation from a standard curve and analyzed by one-way analysis of variance (ANOVA). Cell wall α-1,4-glucan was measured by the same method, but using 1.25 µL of AI resuspension (see below) per reaction.

### Freeze substitution electron microscopy and Thiery staining

Strains were grown for 6 hours in 5 mL YPD at 37°C, then subcultured into 25 mL SD+Gal and grown for a further 18 hours at 37°C. Cells were resuspended in 20% bovine serum albumin/PBS as a cryoprotectant and placed in specimen planchettes, which were high-pressure frozen in a Leica EM PACT2 high-pressure freezer (Leica Microsystems) at −180°C, 2,100 bar and maintained under liquid nitrogen. Samples were then trans-ferred to freeze substitution medium (acetone containing 4% osmium tetroxide (Ted Pella)) under liquid ni-trogen and placed in the Leica AFS automatic freeze substitution system (Leica Microsystems), precooled to −130°C. For freeze substitution, samples were brought to −90°C over a period of 1 h, kept at −90°C for 10 h and subsequently warmed to −20°C over a period of 18 hr. Samples were placed at 4°C for 30 min, washed with anhydrous acetone at room temperature, and then infiltrated and embedded in Eponate 12 resin (Ted Pella).

Sections of 95 nm were cut with a Leica Ultracut UCT ultramicrotome (Leica Microsystems) and placed on freshly glow-discharged nickel grids. Glycogen was detected histochemically by the periodic acid thiocarbo-hydrazide silver proteinate method, modified from Thiery^63,74^. All grid washing steps below were performed by floating each grid in 2 mL dH_2_O in one well of a 24-well plate placed on a rocker. Nickel grids containing ultrathin sections were treated with 1% periodic acid in dH_2_O for 20 min at room temperature and washed 3x 10 min in dH_2_O. Grids were then placed in 1% thiocarbohydrazide/20% acetic acid for 2 min at 60°C pro-tected from light. Grids were rinsed 3x 10 min in dH_2_O, placed in 0.1% silver proteinate in dH_2_O for 5 min at 60°C in the dark, and then washed with dH_2_O 3x 10 min. Samples were viewed on a JEOL 1200 EX transmis-sion electron microscope (JEOL USA) equipped with an AMT 8-megapixel digital camera (Advanced Micros-copy Techniques).

Controls included samples that were histochemically labeled without the initial step of periodic oxidation and nonoxidized sections treated with thiocarbohydrizide and silver proteinate as described above. None of these showed specific silver deposition.

### Quantitation of EM images

To quantify the density of glycogen rosettes at the cytosolic periphery, we chose 10 representative EM images per strain, excluding those with major artifacts. For each image, we measured 500 nm into the cell from the plasma membrane (BioRender). Any cytosol not included in that 500 nm band was blocked out and the area of visible cytosol was measured using ImageJ. Three volunteers then counted the rosettes in each processed image, all of which were deidentified. Reliability among counters was excellent (intraclass correlation coefficient = 0.9153) so we averaged the three counts. Density was calculated as number of rosettes/µm^2^.

### Biotin labeling

*Cn* was inoculated and grown for 6 h in 5 mL YPD at 30°C, then sub-cultured into 25 mL SD+Gal and grown for a further 18 h at 30°C. Four such cultures were combined, spun (3,000 rcf; 5 min; 4°C), and washed 3x with 10 mL cold PBS. Aliquots of 5 x 10^8^ cells were sedimented in microfuge tubes and resuspended in a total of 500 µL cold PBS. Some samples were treated with the indicated amounts of Triton X-100 and rotated for 1 h at 4°C prior to biotin labeling; samples without TX-100 were kept on ice. To biotinylate intact cells (“int + B”), 150 µL freshly prepared No-Weigh Sulfo-NHS-LC-LC-Biotin (Thermo Scientific A35358) was added and the samples were incubated, rotating, for 30 min at 4°C before centrifugation (16,000 rcf; 10 min; 4°C). Superna-tant fractions were reserved, and the pellets were washed 3x with 1 mL cold PBS containing 100 mM glycine to quench any remaining biotin. Control cells (“no B”) received no biotin or washes and were simply collected by centrifugation. All samples were then lysed before SDS-PAGE analysis. For this, pellets were resuspended in 600 µL PBS containing 1X protease inhibitor cocktail (PIC, Sigma P8215) and 1 µL benzonase, transferred to a screw cap tube containing 500 µL 0.5 mm silica beads (Thomas Scientific), subjected to four rounds of bead-beating (each 1 min followed by a 1 min incubation on ice); 500 µL lysate was then immediately transferred to a new tube. For biotinylation of already broken cells (“lys + B”), lysates of unlabeled cells were mixed with 150 µL freshly prepared No-Weigh Sulfo-NHS-LC-LC-Biotin and incubated for 2 h, rotating, at 4°C. A commer-cial bicinchoninic acid assay (Pierce 23225) was used to quantify the total protein in each sample.

### Antibody generation and immunoblot

Affinity-purified polyclonal rabbit antibody to a His-tagged recombinant Goe1 peptide (amino acids 91 to 366) expressed in *E. coli* was prepared by GenScript (NJ, USA). Samples (lysate containing 25 µg total protein or 16 µL supernatant fraction prepared as described in the previous section) were resolved on 15% SDS-PAGE gels, transferred to nitrocellulose, and probed with α-Goe1 antibody (diluted 1:400 in TBS-T containing 1% milk) followed by Goat anti-Rabbit Alexa Fluor 488 (1:10000 dilution in TBS-T) before imaging using a Li-Cor Odyssey instrument.

### Cell wall preparation and assay

Strains were grown for 6 h in 5 mL YPD at 37°C, then subcultured into 25 mL SD+Gal and grown for an addi-tional 42 h shaking at 37°C. To isolate the alkali-insoluble (AI) fraction, we followed the protocol of Aklujkar *et al*^57^. Briefly, 4M KOH was added to each washed pellet (2 mL/g wet weight) and the samples heated at 100°C for 1 h. After neutralization with HCl, samples were centrifuged (3,220 rcf; 30 min; RT), resulting in a gelatinous AI pellet. Of note, the *goe1*Δ pellets were whiter and flakier than those of the other strains. The AI fraction was then washed with 1 mL water until the resulting supernatant did not react with iodine when spotted onto filter paper (usually 3-5 washes with 30-min spins). The supernatant and wash fractions were combined to generate the alkali-soluble (AS) fraction. Both fractions were lyophilized and resuspended in water (2 mL/g original wet weight).

### 1D NMR spectroscopy

Three alkali-insoluble samples for each strain (biological replicates) were combined and re-lyophilized to gen-erate enough material for NMR and linkage analysis (below). Each combined sample was resuspended in 2 mL of water, exhaustively dialyzed (1 kD cutoff) for 72 hours against dH_2_O, transferred to a new tube, and lyophilized.

A portion of 3.7-4.2 mg of each sample was weighed and suspended in 600 µL DMSO-d_6_/pyridinium chloride-d_6_ (1%, w/v). Samples were heated at 40°C for 2 h and each supernatant transferred into a 5-mm NMR tube. Liquid ^1^H-NMR data were obtained at 50 °C on a Bruker Neo 600 MHz spectrometer (1H, 599.66 MHz) with ^1^H-NMR parameters as follows: 60 s relaxation delay, 8992.8 Hz spectral width, 15323 data points and 32 transients with total recycle delay of 2.7 s between each transient. Prior to the Fourier transformation, the data were apodized with an exponential decay function with line broadening of 0.5 Hz, 90⁰ sine square, and zero-filled to 64k points. The baselines were corrected automatically by subtracting a 3rd-order polynomial. The spectra were processed and analyzed with MestreNova (version ×64).

### Glycosyl linkage analysis

Glycosyl linkage analysis was performed by combined gas chromatography-mass spectrometry (GC-MS) of partially methylated alditol acetate (PMAA) derivatives produced from the samples, in a procedure slightly modified from Heiss *et al*^75^. Lyophilized samples (150-290 μg) were first suspended in 300 µL of anhydrous dimethyl sulfoxide (DMSO) and the contents stirred over 3 days. Permethylation was achieved by four rounds of treatment with sodium hydroxide (NaOH) suspension (prepared as in Anumula and Taylor^76^) and iodome-thane (CH_3_I), as follows: first, 300 µL of NaOH suspension was added to each sample and the mixture was magnetically stirred for 15 min at RT. Then, 70 µL CH_3_I was added and the sample stirred for an additional 20 min before a second round of base (15 min) and then CH_3_I (25 min) were added. Each sample was next dis-solved in dichloromethane (DCM) and washed 5 times with 2 mL of water. The water was removed, remaining DCM dried off under a stream of nitrogen, and the sample lyophilized, after which a third and fourth round of base (15 min) and CH_3_I (25 min) treatment were performed. Finally, the sample was washed with DCM and water 5 times, the water was removed, the remaining DCM was dried off under a stream of nitrogen, and the residue was lyophilized.

The permethylated materials were hydrolyzed with 2 M trifluoroacetic acid (TFA) for 2 h at 121°C and dried with isopropanol under a stream of nitrogen. The samples were then reduced with NaBD_4_ in nanopure water overnight, neutralized with glacial acetic acid, and dried with methanol. Finally, the samples were *O*-acety-lated by the addition of 250 µL acetic anhydride and 250 µL TFA at 50°C for 20 min, dried under a stream of nitrogen, reconstituted in DCM, and washed with nanopure water before injection into GC-MS. The PMAAs were analyzed on an Agilent 7890A GC interfaced to a 5975C MSD and separation was performed on a Supelco 2331 fused silica capillary column (30 m x 0.25 mm ID) with a temperature gradient detailed in Table S2.

### β-1,3-glucan assay

Similar to the glycogen assay described above, we measured β-1,3-glucan with an assay that first specifically hydrolyzes β-1,3-glucan and then detects liberated glucose with a fluorescent probe (abcam 303728). 1.25 µL of AI or AS material was used per reaction and all reactions were performed in triplicate. A fourth reaction per sample was performed in parallel without enzyme as a background control; free glucose was never de-tected in these samples. Fluorescence intensity was measured using a Biotek plate reader reading excita-tion/emission = 535/587 in end-point mode at 37°; gain was set to 60 V. Background (intensity of the 0 µg standard) was subtracted from each sample and β-1,3-glucan standard. β-1,3-glucan content of samples was calculated from interpolation of the standard curve and compared by one-way ANOVA.

### Intracellular survival assay

As previously described^77^, THP-1 cells were differentiated by 48 h incubation in medium containing 25 nM phorbol myristate acetate (PMA) in 24-well plates (1.7 x 10^5^ cells/well) and allowed to recover by 24 h in medium without PMA. In parallel, fungal cells were grown overnight in YPD, washed with sterile PBS, and adjusted to a 10^7^ cells/ml in PBS before being opsonized in 40% human serum in PBS (30 min; 37°C). Opsonized fungi were washed, resuspended in RPMI, and added to differentiated cells at a multiplicity of infection of 0.1 in triplicate wells of three parallel plates. After 1 h of incubation at 37°C and 5% CO_2_, all plates were washed with PBS. Distilled water was added to one plate to lyse the THP-1 cells and the resulting lysate was immediately plated on YPD (time 0). The remaining plates were refilled with THP-1 medium and incubated for 24 or 48 h prior to THP-1 cell lysis and plating. Resulting colony forming units (CFU) were enumerated and compared using two-way ANOVA.

### Animal studies

Cryptococcal strains were grown overnight in 5 mL YPD, washed, and diluted in sterile PBS to 2.5 x 10^5^ cells/ml. Female 6-week-old C57BL/6 mice (Jackson Laboratory) were anesthetized by subcutaneous injection of 120 µl of 10 mg/mL ketamine and 2 mg/ml xylazine in sterile water and intranasally inoculated with 1.25 x 10^4^ cells in 25 µl PBS. Mice were humanely sacrificed 9 days after infection and homogenates prepared from harvested lungs were plated on YPD. Resulting CFU were enumerated and analyzed by one-way ANOVA with Tukey’s *post hoc* test.

### Ethics statement

All animal studies were approved by the Washington University Institutional Animal Care and Use Committee (Protocol #20-0108) and conducted according to the “Guide for the Care and Use of Laboratory Animals” published by the National Research Council and endorsed by the Association for the Assessment and Accred-itation of Laboratory Animal Care.

## ACKNOWLEDGEMENTS

Glycan NMR and linkage analysis was performed by Christian Heiss, Li Tan, and Jie Lun Cheng at the Complex Carbohydrate Research Center, which is supported by the U.S. Department of Energy, Office of Science, Basic Energy Sciences, Chemical Sciences, Geosciences and Biosciences Division, under award #DE-SC0015662. The *gsy1*Δ and *glc3*Δ strains used in this study were generated as part of a deletion library funded by R01 AI100272 to the Madhani lab and purchased from the Fungal Genetics Stock Center. Special thanks to Wandy Beatty in the Molecular Microbiology Imaging Facility, who generated the beautiful EM images in this paper. We are also grateful to Camaron Hole for pointing us to the literature on cell wall α-1,4-glucan; Mario Feldman for discussions about protein topology; Thomas Hurtaux for his insights on glycosyltransferases; Elizabeth Gaylord, Daphne Ko, and John Loza for helpful comments on the manuscript and figures; John Loza, Caroline Stone-Viau, and Max Stone-Viau for their time counting glycogen rosettes (an extra thanks to C. Stone-Viau for her statistical expertise); Shannon Kuo for general lab assistance; and Wayne Wilson for generously providing *Sc* strains. All schematics were created using BioRender.

## Author contributions

Conceptualization, L.L. and T.L.D; Methodology, L.L. and T.L.D; Investigation, Formal Analysis, Writing – Original Draft, Visualization, L.L; Resources, Supervision, Writing – Review & Editing, Project Administration, T.L.D; Funding Acquisition, L.L. and T.L.D.

## Declaration of interests

The authors declare no competing interests.

## Funding

Support for this project was provided by National Institute of Allergy and Infectious Diseases awards R01 135012 and R21 175875 to T.L.D. and F31 150194 to L.L.

## Notes

### Competing Interest Statement

The authors have declared no competing interest.

## REFERENCES

1. Wilson, W.A., Roach, P.J., Montero, M., Baroja-Fernández, E., Muñoz, F.J., Eydallin, G., Viale, A.M., and Pozueta-Romero, J. (2010). Regulation of glycogen metabolism in yeast and bacteria. FEMS Microbiol Rev 34, 952–985. 10.1111/j.1574-6976.2010.00220.x.

2. Shearer, J., and Graham, T.E. (2004). Novel Aspects of Skeletal Muscle Glycogen and Its Regulation Dur-ing Rest and Exercise. Exercise and Sport Sciences Reviews 32, 120–126.

3. Mundkur, B. (1960). Electron microscopical studies of frozen-dried yeast: I. Localization of polysaccha-rides. Experimental Cell Research 20, 28–42. 10.1016/0014-4827(60)90219-6.

4. Cheng, C., Mu, J., Farkas, I., Huang, D., Goebl, M.G., and Roach, P.J. (1995). Requirement of the self-glu-cosylating initiator proteins Glg1p and Glg2p for glycogen accumulation in *Saccharomyces cerevisiae*. Mol Cell Biol 15, 6632–6640.

5. Mu, J., Cheng, C., and Roach, P.J. (1996). Initiation of glycogen synthesis in yeast: requirement of multi-ple tyrosine residues for function of the self-glucosylating Glg proteins *in vivo*. J. Biol. Chem. 271, 26554–26560. 10.1074/jbc.271.43.26554.

6. Farkas, I., Hardy, A., DePaoli-Roachll, A., and Roach, J. Isolation of the GSY1 gene encoding yeast glyco-gen synthase and evidence for the existence of a second gene. 9.

7. Farkas, I., Hardy, T.A., Goebl, M.G., and Roach, P.J. (1991). Two glycogen synthase isoforms in *Saccharo-myces cerevisiae* are coded by distinct genes that are differentially controlled. J. Biol. Chem. 266, 15602–15607.

8. Gunja, Z.H., Manners, D.J., and Maung, K. (1960). Studies on carbohydrate-metabolizing enzymes. 3. Yeast branching enzyme. Biochem J 75, 441–450.

9. Manners, D.J. (1971). The structure and biosynthesis of storage carbohydrates in yeast. In The Yeasts, A. Rose and J. Harrison, eds. (Academic Press), pp. 419–439.

10. Hwang, P.K., Tugendreich, S., and Fletterick, R.J. (1989). Molecular analysis of *GPH1*, the gene encoding glycogen phosphorylase in *Saccharomyces cerevisiae*. Mol Cell Biol 9, 1659–1666. 10.1128/mcb.9.4.1659-1666.1989.

11. Rath, V.L., Hwang, P.K., and Fletterick, R.J. (1992). Purification and crystallization of glycogen phosphory-lase from *Saccharomyces cerevisiae*. Journal of Molecular Biology 225, 1027–1034. 10.1016/0022-2836(92)90102-P.

12. Teste, M.A., Enjalbert, B., Parrou, J.L., and François, J.M. (2000). The *Saccharomyces cerevisiae YPR184w* gene encodes the glycogen debranching enzyme. FEMS Microbiol Lett 193, 105–110. 10.1111/j.1574-6968.2000.tb09410.x.

13. Rajasingham, R., Govender, N.P., Jordan, A., Loyse, A., Shroufi, A., Denning, D.W., Meya, D.B., Chiller, T.M., and Boulware, D.R. (2022). The global burden of HIV-associated cryptococcal infection in adults in 2020: a modelling analysis. The Lancet Infectious Diseases 22, 1748–1755. 10.1016/S1473-3099(22)00499-6.

14. Levitz, S.M. (1991). The ecology of *Cryptococcus neoformans* and the epidemiology of cryptococcosis. Reviews of Infectious Diseases 13, 1163–1169.

15. Nielsen, K., De Obaldia, A.L., and Heitman, J. (2007). *Cryptococcus neoformans* mates on pigeon guano: implications for the realized ecological niche and globalization. Eukaryot Cell 6, 949–959. 10.1128/EC.00097-07.

16. Kamari, A., Sepahvand, A., and Mohammadi, R. (2017). Isolation and molecular characterization of *Cryptococcus* species isolated from pigeon nests and Eucalyptus trees. Curr Med Mycol 3, 20–25. 10.29252/cmm.3.2.20.

17. Gaylord, E.A., Choy, H.L., and Doering, T.L. (2020). Dangerous liaisons: interactions of *Cryptococcus neoformans* with host phagocytes. Pathogens 9. 10.3390/pathogens9110891.

18. Tseng, H.-K., Huang, T.-Y., Wu, A.Y.-J., Chen, H.-H., Liu, C.-P., and Jong, A. (2015). How *Cryptococcus* interacts with the blood-brain barrier. Future Microbiol 10, 1669–1682. 10.2217/fmb.15.83.

19. Santiago-Tirado, F.H., Onken, M.D., Cooper, J.A., Klein, R.S., and Doering, T.L. (2017). Trojan horse transit contributes to blood-brain barrier crossing of a eukaryotic pathogen. mBio 8. 10.1128/mBio.02183-16.

20. Esher, S.K., Zaragoza, O., and Alspaugh, J.A. (2018). Cryptococcal pathogenic mechanisms: a dangerous trip from the environment to the brain. Mem Inst Oswaldo Cruz 113, e180057. 10.1590/0074-02760180057.

21. Philips, B.J., Meguer, J.-X., Redman, J., and Baker, E.H. (2003). Factors determining the appearance of glucose in upper and lower respiratory tract secretions. Intensive Care Med 29, 2204–2210. 10.1007/s00134-003-1961-2.

22. Baker, E.H., Clark, N., Brennan, A.L., Fisher, D.A., Gyi, K.M., Hodson, M.E., Philips, B.J., Baines, D.L., and Wood, D.M. (2007). Hyperglycemia and cystic fibrosis alter respiratory fluid glucose concentrations estimated by breath condensate analysis. Journal of Applied Physiology 102, 1969–1975. 10.1152/jap-plphysiol.01425.2006.

23. Gurung, P., Zubair, M., and Jialal, I. (2023). Plasma Glucose. In StatPearls (StatPearls Publishing).

24. Irani, D.N. (2009). Properties and composition of normal cerebrospinal fluid. In Cerebrospinal Fluid in Clinical Practice, D. N. Irani, ed. (W.B. Saunders), pp. 69–89. 10.1016/B978-141602908-3.50013-3.

25. Petzold, E.W., Himmelreich, U., Mylonakis, E., Rude, T., Toffaletti, D., Cox, G.M., Miller, J.L., and Perfect, J.R. (2006). Characterization and regulation of the trehalose synthesis pathway and Its importance in the pathogenicity of *Cryptococcus neoformans*. Infect Immun 74, 5877–5887. 10.1128/IAI.00624-06.

26. Botts, M.R., Huang, M., Borchardt, R.K., and Hull, C.M. (2014). Developmental cell fate and virulence are linked to trehalose homeostasis in *Cryptococcus neoformans*. Eukaryot Cell 13, 1158–1168. 10.1128/EC.00152-14.

27. Ortiz, S.C., McKeon, M.C., Botts, M.R., Gage, H., Frerichs, A., and Hull, C.M. (2023). Spores of the fungal pathogen *Cryptococcus* exhibit cell type-specific carbon source utilization during germination. Preprint at bioRxiv, 10.1101/2023.10.01.560341 10.1101/2023.10.01.560341.

28. Lin, X., and Heitman, J. (2005). Chlamydospore formation during hyphal growth in *Cryptococcus neoformans*. Eukaryot Cell 4, 1746–1754. 10.1128/EC.4.10.1746-1754.2005.

29. Banks, I.R., Specht, C.A., Donlin, M.J., Gerik, K.J., Levitz, S.M., and Lodge, J.K. (2005). A chitin synthase and its regulator protein are critical for chitosan production and growth of the fungal pathogen *Crypto-coccus neoformans*. Eukaryot Cell 4, 1902–1912. 10.1128/EC.4.11.1902-1912.2005.

30. Baker, L.G., Specht, C.A., Donlin, M.J., and Lodge, J.K. (2007). Chitosan, the deacetylated form of chitin, is necessary for cell wall integrity in *Cryptococcus neoformans*. Eukaryot Cell 6, 855–867. 10.1128/EC.00399-06.

31. Thompson, J.R., Douglas, C.M., Li, W., Jue, C.K., Pramanik, B., Yuan, X., Rude, T.H., Toffaletti, D.L., Perfect, J.R., and Kurtz, M. (1999). A glucan synthase *FKS1* homolog in *Cryptococcus neoformans* is single copy and encodes an essential function. J Bacteriol 181, 444–453.

32. Hu, X., Yang, P., Chai, C., Liu, J., Sun, H., Wu, Y., Zhang, M., Zhang, M., Liu, X., and Yu, H. (2023). Struc-tural and mechanistic insights into fungal β-1,3-glucan synthase FKS1. Nature 616, 190–198. 10.1038/s41586-023-05856-5.

33. Beattie, S.R., Jezewski, A.J., Ristow, L.C., Wellington, M., and Krysan, D.J. (2022). *FKS1* is required for *Cryptococcus neoformans* fitness *in vivo*: application of copper-regulated gene expression to mouse models of cryptococcosis. mSphere 7, e0016322. 10.1128/msphere.00163-22.

34. Kalem, M.C., Subbiah, H., Leipheimer, J., Glazier, V.E., and Panepinto, J.C. (2021). Puf4 mediates post-transcriptional regulation of cell wall biosynthesis and caspofungin resistance in *Cryptococcus neoformans*. mBio 12, 10.1128/mbio.03225-20. 10.1128/mbio.03225-20.

35. Johnson, M.D., and Perfect, J.R. (2003). Caspofungin: first approved agent in a new class of antifungals. Expert Opin Pharmacother 4, 807–823. 10.1517/14656566.4.5.807.

36. Reese, A.J., and Doering, T.L. (2003). Cell wall α-1,3-glucan is required to anchor the *Cryptococcus neoformans* capsule. Mol Microbiol 50, 1401–1409. 10.1046/j.1365-2958.2003.03780.x.

37. Reese, A.J., Yoneda, A., Breger, J.A., Beauvais, A., Liu, H., Griffith, C.L., Bose, I., Kim, M.-J., Skau, C., Yang, S., et al. (2007). Loss of cell wall alpha(1-3) glucan affects *Cryptococcus neoformans* from ultrastructure to virulence. Mol Microbiol 63, 1385–1398. 10.1111/j.1365-2958.2006.05551.x.

38. Cabib, E., Blanco, N., Grau, C., Rodríguez-Peña, J.M., and Arroyo, J. (2007). Crh1p and Crh2p are required for the cross-linking of chitin to β(1-6)glucan in the *Saccharomyces cerevisiae* cell wall. Molecular Micro-biology 63, 921–935. 10.1111/j.1365-2958.2006.05565.x.

39. Montijn, R.C., Vink, E., Müller, W.H., Verkleij, A.J., Van Den Ende, H., Henrissat, B., and Klis, F.M. (1999). Localization of synthesis of β1,6-glucan in *Saccharomyces cerevisiae*. Journal of Bacteriology 181, 7414– 7420. 10.1128/jb.181.24.7414-7420.1999.

40. Boone, C., Sommer, S.S., Hensel, A., and Bussey, H. (1990). Yeast *KRE* genes provide evidence for a path-way of cell wall beta-glucan assembly. J Cell Biol 110, 1833–1843.

41. Roemer, T., Delaney, S., and Bussey, H. (1993). *SKN1* and *KRE6* define a pair of functional homologs en-coding putative membrane proteins involved in beta-glucan synthesis. Mol Cell Biol 13, 4039–4048.

42. Roemer, T., Paravicini, Gerhard, Payton, Mark A., and Bussey, Howard (1994). Characterization of the yeast (1-->6)-beta-glucan biosynthetic components, Kre6p and Skn1p, and genetic interactions between the *PKC1* pathway and extracellular matrix assembly. J Cell Biol 127, 567–579.

43. Brown, J.L., Kossaczka, Z., Jiang, B., and Bussey, H. (1993). A mutational analysis of killer toxin resistance in *Saccharomyces cerevisiae* identifies new genes involved in cell wall (1-> 6)-β-glucan synthesis. Genet-ics 133, 837–849.

44. Sacher, M., Barrowman, J., Schieltz, D., Yates, J.R., and Ferro-Novick, S. (2000). Identification and charac-terization of five new subunits of TRAPP. European Journal of Cell Biology 79, 71–80. 10.1078/S0171-9335(04)70009-6.

45. Breinig, F., Schleinkofer, K., and Schmitt, M.J. (2004). Yeast Kre1p is GPI-anchored and involved in both cell wall assembly and architecture. Microbiology (Reading) 150, 3209–3218. 10.1099/mic.0.27175-0.

46. Brown, J.L., and Bussey, H. (1993). The yeast *KRE9* gene encodes an O glycoprotein involved in cell sur-face beta-glucan assembly. Mol Cell Biol 13, 6346–6356.

47. Dijkgraaf, G.J.P., Brown, J.L., and Bussey, H. (1996). The *KNH1* gene of *Saccharomyces cerevisiae* is a functional homolog of KRE9. Yeast 12, 683–692. 10.1002/(SICI)1097-0061(19960615)12:7<683::AID-YEA959>3.0.CO;2-8.

48. Klebl, F., and Tanner, W. (1989). Molecular cloning of a cell wall exo-beta-1,3-glucanase from *Saccharo-myces cerevisiae*. J Bacteriol 171, 6259–6264.

49. Fankhauser, C., and Conzelmann, A. (1991). Purification, biosynthesis and cellular localization of a major 125-kDa glycophosphatidylinositol-anchored membrane glycoprotein of *Saccharomyces cerevisiae*. Eu-ropean Journal of Biochemistry 195, 439–448. 10.1111/j.1432-1033.1991.tb15723.x.

50. Aimanianda, V., Simenel, C., Garnaud, C., Clavaud, C., Tada, R., Barbin, L., Mouyna, I., Heddergott, C., Popolo, L., Ohya, Y., et al. (2017). The dual activity responsible for the elongation and branching of β-(1,3)-glucan in the fungal cell wall. mBio 8, 10.1128/mbio.00619-17. 10.1128/mbio.00619-17.

51. Gilbert, N.M., Donlin, M.J., Gerik, K.J., Specht, C.A., Djordjevic, J.T., Wilson, C.F., Sorrell, T.C., and Lodge, J.K. (2010). *KRE* genes are required for beta-1,6-glucan synthesis, maintenance of capsule architecture and cell wall protein anchoring in *Cryptococcus neoformans*. Mol Microbiol 76, 517–534. 10.1111/j.1365-2958.2010.07119.x.

52. Ling, A.R., Nanji, D.R., and Paton, F.J. (1925). Studies on glycogen, Part I.-The nature of yeast glycogen, its preparation, estimation, and its role in yeast metabolism. Journal of the Institute of Brewing 31, 316– 321. 10.1002/j.2050-0416.1925.tb04919.x.

53. Trevelyan, W.E., and Harrison, J.S. (1952). Studies on yeast metabolism. 1. Fractionation and microdeter-mination of cell carbohydrates. Biochem J 50, 298–303.

54. Gunja-Smith, Z., and Smith, E.E. (1974). Evidence for the periplasmic location of glycogen in *Saccharomyces*. Biochemical and Biophysical Research Communications 56, 588–592. 10.1016/0006-291X(74)90644-5.

55. Gunja-Smith, Z., Patil, N.B., and Smith, E.E. (1977). Two pools of glycogen in *Saccharomyces*. J Bacteriol 130, 818–825.

56. Arvindekar, A.U., and Patil, N.B. (2002). Glycogen--a covalently linked component of the cell wall in *Saccharomyces cerevisiae*. Yeast 19, 131–139. 10.1002/yea.802.

57. Aklujkar, P.P., Sankh, S.N., and Arvindekar, A.U. (2008). A simplified method for the isolation and estimation of cell wall bound glycogen in *Saccharomyces cerevisiae*. Journal of the Institute of Brewing 114, 205–208. 10.1002/j.2050-0416.2008.tb00330.x.

58. Lowman, D.W., Sameer Al-Abdul-Wahid, M., Ma, Z., Kruppa, M.D., Rustchenko, E., and Williams, D.L. (2021). Glucan and glycogen exist as a covalently linked macromolecular complex in the cell wall of *Candida albicans* and other *Candida* species. The Cell Surface 7, 100061. 10.1016/j.tcsw.2021.100061.

59. James, P.G., Cherniak, R., Jones, R.G., Stortz, C.A., and Reiss, E. (1990). Cell-wall glucans of *Cryptococcus neoformans* Cap67. Carbohydr Res 198, 23–38. 10.1016/0008-6215(90)84273-w.

60. Notredame, C., Higgins, D.G., and Heringa, J. (2000). T-Coffee: a novel method for fast and accurate mul-tiple sequence alignment. Journal of Molecular Biology 302, 205–217. 10.1006/jmbi.2000.4042.

61. Lin, A., Mu, J., Yang, J., and Roach, P.J. (1999). Self-glucosylation of glycogenin, the initiator of glycogen biosynthesis, involves an inter-subunit reaction. Archives of Biochemistry and Biophysics 363, 163–170. 10.1006/abbi.1998.1073.

62. Pavelka, M., and Roth, J. (2010). Glycogen. In Functional Ultrastructure: Atlas of Tissue Biology and Pathology, M. Pavelka and J. Roth, eds. (Springer), pp. 140–140. 10.1007/978-3-211-99390-3_73.

63. Coulary, B., Aigle, M., and Schaeffer, J. (2001). Evidence for glycogen structures associated with plasma membrane invaginations as visualized by freeze-substitution and the Thiery reaction in *Saccharomyces cerevisiae*. Journal of Electron Microscopy 50, 133–137. 10.1093/jmicro/50.2.133.

64. Terashima, M., Kim, K.M., Adachi, T., Nielsen, P.J., Reth, M., Köhler, G., and Lamers, M.C. (1994). The IgM antigen receptor of B lymphocytes is associated with prohibitin and a prohibitin-related protein. EMBO J 13, 3782–3792.

65. Nazari, M., Kurdi, M., and Heerklotz, H. (2012). Classifying surfactants with respect to their effect on lipid membrane order. Biophys J 102, 498–506. 10.1016/j.bpj.2011.12.029.

66. Fan, W., Kraus, P.R., Boily, M.-J., and Heitman, J. (2005). *Cryptococcus neoformans* gene expression during murine macrophage infection. Eukaryotic Cell 4, 1420–1433. 10.1128/ec.4.8.1420-1433.2005.

67. Torija, M.-J., Novo, M., Lemassu, A., Wilson, W., Roach, P.J., François, J., and Parrou, J.-L. (2005). Glycogen synthesis in the absence of glycogenin in the yeast *Saccharomyces cerevisiae*. FEBS Letters 579, 3999–4004. 10.1016/j.febslet.2005.06.007.

68. Fonzi, W.A. (1999). *PHR1* and *PHR2* of *Candida albicans* encode putative glycosidases required for proper cross-linking of beta-1,3- and beta-1,6-glucans. J Bacteriol 181, 7070–7079. 10.1128/JB.181.22.7070-7079.1999.

69. Fu, J., Hettler, E., and Wickes, B.L. (2006). Split marker transformation increases homologous integration frequency in *Cryptococcus neoformans*. Fungal Genet. Biol. 43, 200–212. 10.1016/j.fgb.2005.09.007.

70. Kwon-Chung, K.J., Edman, J.C., and Wickes, B.L. (1992). Genetic association of mating types and viru-lence in *Cryptococcus neoformans*. Infect Immun 60, 602–605.

71. Lengeler, K.B., Fox, D.S., Fraser, J.A., Allen, A., Forrester, K., Dietrich, F.S., and Heitman, J. (2002). Mating-type locus of *Cryptococcus neoformans*: a step in the evolution of sex chromosomes. Eukaryot Cell 1, 704–718. 10.1128/EC.1.5.704-718.2002.

72. Nielsen, K., Cox, G.M., Wang, P., Toffaletti, D.L., Perfect, J.R., and Heitman, J. (2003). Sexual Cycle of *Cryptococcus neoformans* var. *grubii* and Virulence of Congenic a and α Isolates. Infect Immun 71, 4831– 4841. 10.1128/IAI.71.9.4831-4841.2003.

73. Gietz, R.D., and Schiestl, R.H. (2007). Quick and easy yeast transformation using the LiAc/SS carrier DNA/PEG method. Nat Protoc 2, 35–37. 10.1038/nprot.2007.14.

74. Thiery, J.P. (1967). Mise en évidence des polysaccharides sur coupes fines en microscopie électronique. J Microscopie 6, 987–1018.

75. Heiss, C., Klutts, J.S., Wang, Z., Doering, T.L., and Azadi, P. (2009). The structure of *Cryptococcus neofor-mans* galactoxylomannan contains β-D-glucuronic acid. Carbohydr Res 344, 915–920. 10.1016/j.carres.2009.03.00.

76. Anumula, K.R., and Taylor, P.B. (1992). A comprehensive procedure for preparation of partially methyl-ated alditol acetates from glycoprotein carbohydrates. Analytical Biochemistry 203, 101–108. 10.1016/0003-2697(92)90048-C.

77. Chang, A.L., and Doering, T.L. (2018). Maintenance of Mitochondrial Morphology in *Cryptococcus neoformans* Is Critical for Stress Resistance and Virulence. mBio 9, e01375–18, /mbio/9/6/mBio.01375-18.atom. 10.1128/mBio.01375-18.

