## Supplemental data for "A novel glycosyltransferase organizes fungal glycogen and cell wall glucans"

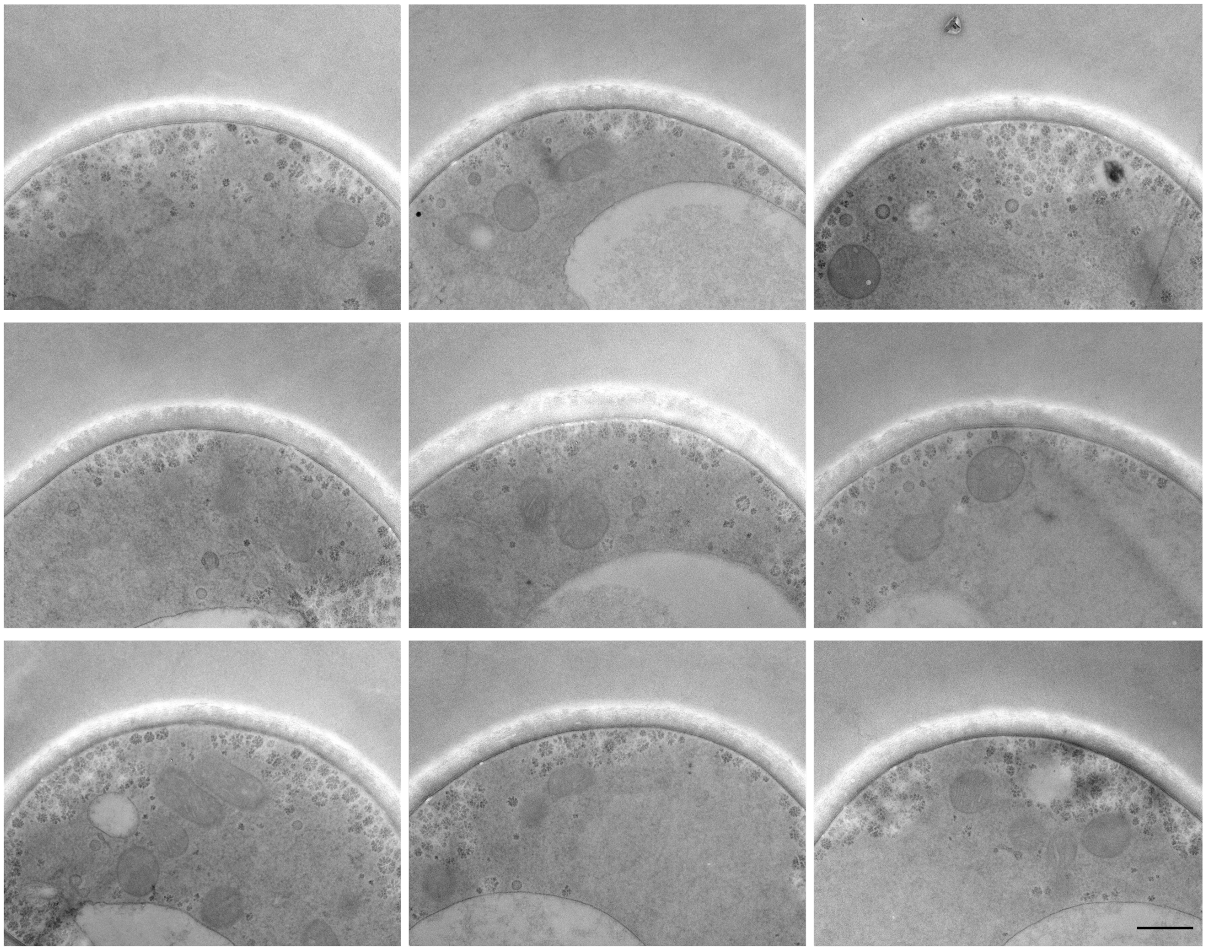

**Figure S1. Electron micrographs of Thiery-stained WT cells.** All images to the same scale; scale bar, 500 nm.

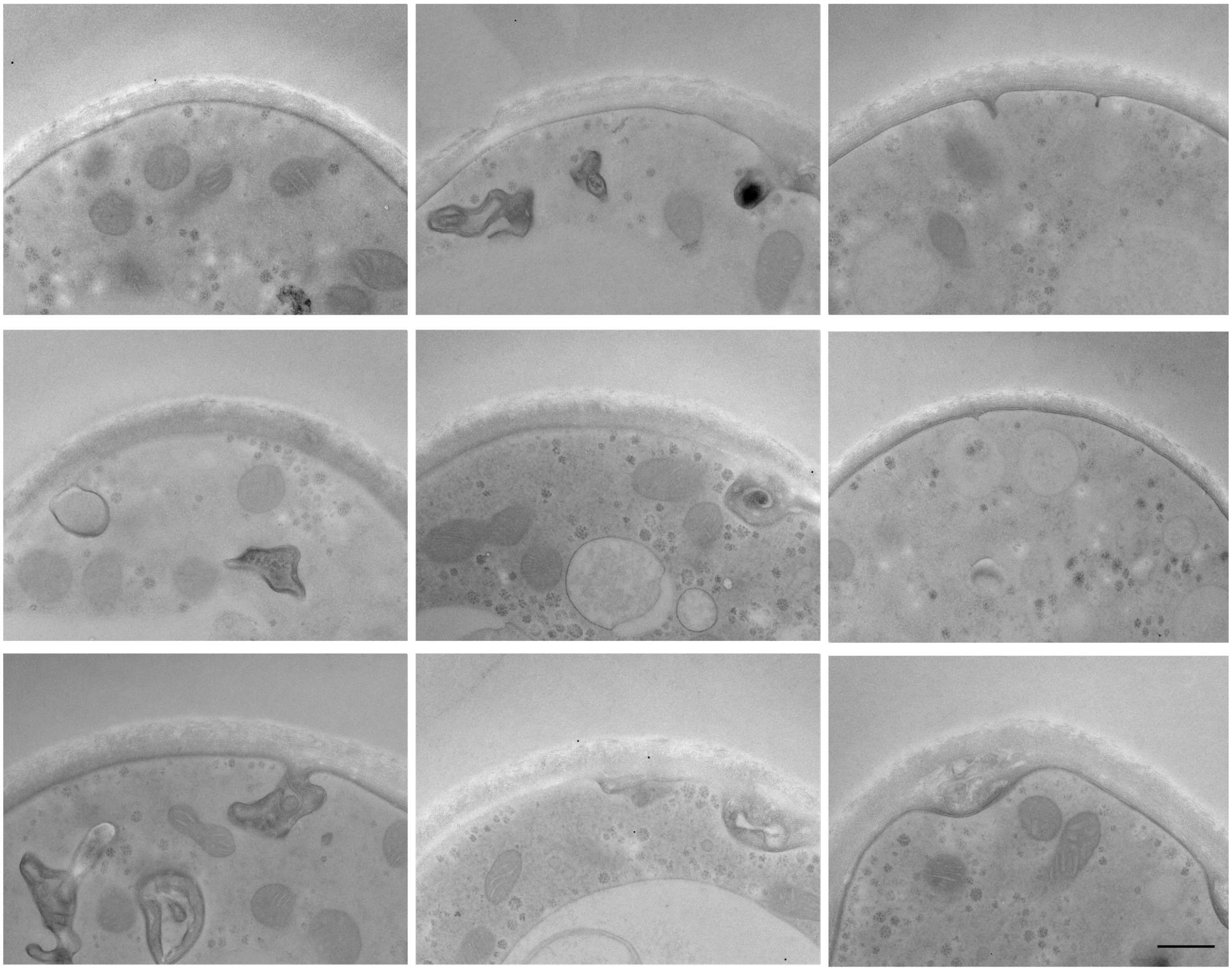

**Figure S2. Electron micrographs of Thiery-stained *goe1*Δ cells.** All images to the same scale; scale bar, 500 nm.

| Sample | WT | *goe1Δ* | *goe1Δ::GOE1* |
|---|---|---|---|
| **Residue** | **% area** | **% area** | **% area** |
| Terminal Glucopyranosyl residue (t-Glc) | 11.02 | 7.16 | 11.88 |
| 3-Linked Glucopyranosyl residue (3-Glc) | 20.12 | 10.56 | 19.10 |
| 6-Linked Glucopyranosyl residue (6-Glc) | 17.62 | 16.36 | 18.48 |
| 4-Linked Glucopyranosyl residue (4-Glc) | 17.64 | 24.07 | 19.72 |
| 3,4-Linked Glucopyranosyl residue (3,4-Glc) | 1.17 | 2.53 | 1.58 |
| 3,6-Linked Glucopyranosyl residue (3,6-Glc) | 10.82 | 6.71 | 10.44 |
| 4,6-Linked Glucopyranosyl residue (4,6-Glc) | 3.03 | 3.94 | 3.14 |
| 2,3,4,6-Linked Glucopyranosyl residue (2,3,4,6-Glc) | 1.04 | 3.81 | 1.43 |

**Table S1. Glycosyl linkage analysis results.** Data shown as relative percentage of each detected linkage. Not shown, species other than glucose or below 1% abundance in WT cells. Note that this method does not distinguish between $\alpha$- and $\beta$-linked residues.

| | Rate (°C/min) | Value (°C) | Hold Time (min) | Run Time (min) |
|---|---|---|---|---|
| Initial | | 60 | 1 | 1 |
| Ramp 1 | 27.5 | 170 | 0 | 5 |
| Ramp 2 | 4 | 235 | 2 | 23.5 |
| Ramp 3 | 3 | 240 | 18 | 42.9 |

**Table S2. Temperature program for the GC-MS analysis of PMAAs.**
